# Maintenance of copy number variation at the human salivary agglutinin gene (*DMBT1*) by balancing selection driven by host-microbe interactions

**DOI:** 10.1101/2021.10.29.466477

**Authors:** Adel F. Alharbi, Nongfei Sheng, Katie Nicol, Nicklas Strömberg, Edward J. Hollox

## Abstract

Most genetic variation in humans occurs in a pattern consistent with neutral evolution, but a small subset is maintained by balancing selection. Identifying loci under balancing selection is important not only for understanding the processes explaining variation in the genome, but also to identify loci with alleles that affect response to the environment and disease. Several genome scans using genetic variation data have identified the 5’ end of the *DMBT1* gene as a region undergoing balancing selection. *DMBT1* encodes the pattern-recognition glycoprotein DMBT1, also known as SALSA, gp340 or salivary agglutinin. It binds to a wide variety of pathogens through a tandemly-arranged scavenger receptor cysteine-rich (SRCR) domain, with the number of SRCR domains varying in humans. Here we use expression analysis, linkage in pedigrees, and long range single transcript sequencing, to show that the signal of balancing selection is driven by one haplotype usually carrying shorter SRCR repeats, and another usually carrying a longer SRCR repeat, within the coding region of *DMBT1*. The DMBT1 protein size isoform encoded by a shorter SRCR domain repeat allele showed complete loss of binding of a cariogenic and invasive *Streptococcus mutans* strain in contrast to the long SRCR allele. Taken together, our results suggest that balancing selection at *DMBT1* is due to host-microbe interactions of encoded SRCR tandem repeat alleles.

## Introduction

Patterns of human genetic variation across populations are consistent with most genetic variation being neutral, with only a small subset of variation affected by natural selection (Fu and Akey, 2013). The role of balancing selection in shaping the pattern of human genetic diversity by maintaining alleles in a population at higher frequencies and/or for long periods of time, remains elusive (Key et al., 2014). From the earliest study on malaria and sickle cell disease to modern genome scans, it is becoming clear that pathogen pressure has been important in maintaining alleles at intermediate frequencies at particular loci. Most evidence has focused on well-established examples, such as the sickle-cell trait protecting against severe malaria in individuals heterozygous for the beta-globin (*HBB*) sickle-cell allele, glucose-6-phosphate dehydrogenase (G6PD), and the major histocompatibility complex (MHC) region (Key et al., 2014; Solberg et al., 2008). In these examples, overdominance (heterozygote advantage) maintains two or more distinct alleles in a population. Other balancing selection mechanisms, such as frequency-dependent selection or fluctuating selection, remain underexplored.

Analysis of polymorphisms maintained in both human and chimpanzee lineages suggest, in addition to the MHC region, that there are 125 further regions of balancing selection enriched for genes likely to be involved in host-pathogen interactions (Leffler et al., 2013). Scans of traditional within-species summary statistics that can detect balancing selection, such as Tajima’s D or HKA tests (Bubb et al., 2006; Andrés et al., 2009), have identified some loci undergoing balancing selection, but miss other well-known examples, suggesting a lack of power of the statistics used. In addition, cryptic duplications or copy number variation (CNV) can be misinterpreted as balancing selection if the genome is not fully or correctly annotated. More accurate annotation of CNV in genomes combined with whole genome sequencing data and statistics designed to sensitively detect haplotypes exhibiting characteristics of balancing selection divergence and frequency have enabled identification of a larger number of balancing selection sites (Bitarello et al., 2018; de Filippo et al., 2016; DeGiorgio et al., 2014; Siewert and Voight, 2017).

In almost all cases where genomewide scans for balancing selection have been done, moving beyond a statistical signal and determining the molecular basis for the balancing selection has been lacking. This is important, not only to provide a clear explanation for patterns of selection and our understanding of the past pressures on humans, but to understand how the variation interacts with pathogens today, with consequences for disease progression, resistance and treatment (Key et al., 2014).

The mucin hybrid glycoprotein DMBT1 (also known as SALSA, gp340, muclin or salivary agglutinin), encoded by the *DMBT1* is a pattern-recognition and scavenger receptor protein (Canton et al., 2013). Accordingly, DMBT1 is composed of tandemly-arranged scavenger receptor cysteine-rich (SRCR) and interspersed glycosylated Ser/Thr-rich motif (SID) domains for microbe and host-ligand binding, and CUB (complement C1r/C1s, Uegf, Bmp1) and ZP (zona pellucida-like) domains for cell polarisation and polymerisation (Mollenhauer et al., 1997; Prakobphol et al., 2000; Reichhardt et al., 2017).

DMBT1 is expressed by macrophages and in tissues encountering microbes, such as saliva, lung, colon, amnionic fluid and meconium. DMBT1 binds many different viruses and bacteria, including *Streptococcus mutans*, and host ligands, such as secretory IgA, lactoferrin and mucin-5B, through the SRCR and SID domains (Thornton et al., 2001; Ligtenberg et al., 2004; Loimaranta et al., 2005). Saliva adhesion and aggregation of streptococcal ligands by DMBT1 and its activation of complement by the lectin pathway have differential outcomes in solution versus on surfaces (Loimaranta et al., 2005; Leito et al., 2011; Reichhardt et al., 2012; Reichhardt and Meri, 2016; Gunput et al., 2016).

The *DMBT1* gene shows extensive multi-allelic copy number variation (CNV) across all populations, with the tandemly-repeated CNV leading to different alleles with between 7-21 SRCR domains (Sasaki et al., 2002; Polley et al., 2015). Saliva DMBT1 protein size isoforms I-IV corresponding in size to shorter and longer isoforms exist (Eriksson et al., 2007; Esberg et al., 2012), and variation in the number of SRCR domains influence binding, with a short isoform (8 SRCR domains) binding bacteria 30-40% less effectively than the longer isoform (13 SRCR domains) (Esberg et al., 2012; Bikker et al., 2017). Multiple transcripts of *DMBT1* have been detected, and are often interpreted as evidence of alternative splicing, although the range and nature of the alternative transcripts are consistent with CNV of the underlying *DMBT1* SRCR exons. The O-glycosylation of the Thr-rich SID domains in DMBT1 by short chain GalGalNAc and long chain type 1 and type 2 Lewis and ABO chains also vary between individuals (Schulz et al., 2002; Eriksson et al., 2007). The Lewis and ABO glycosylation of DMBT1 depends on Secretor (Se) status, encoded by a polymorphism in the fucosyltransferase 2 (*FUT2*) gene (Kelly et al., 1995). However, the relative contributions of variation in SRCR copy number, at the DNA level, alternative splicing at the RNA level, and alternative glycosylation at the post-translational processing level, to the observed variation in DMBT1 protein size between individuals remains to be determined.

The consequences of genetic, translational or post-translational variation on *DMBT1* function and role in disease development remains largely unknown, and DMBT1 complexity, redundancy of pattern recognition pathways and disease heterogeneity may explain difficulties in linking DMBT1 to disease development. Analysis of variation at the population level, combined with the known function of DMBT1 and the binding affinities of the SRCR domains suggested an evolutionary explanation based on an interaction between the human dietary environment and genetic variation affecting affinity to the causative agent of dental caries, the bacteria *Streptococcus mutans* (Polley et al., 2015). This was subsequently supported by direct evidence that different length alleles have differing binding affinities to *S. mutans* (Bikker et al., 2017).

Several genomewide selection scans have identified *DMBT1* as a gene showing one of the strongest signals for balancing selection in Europeans and Africans (Figure 1). Using T1 and T2 tests, which measure the ratio of within-species to between-species variation and any excess of intermediate frequency variants respectively, *DMBT1* was found in the top 20 of genomic loci in both European-Americans from Utah (CEU) and Yoruba from Ibadan in Nigeria (YRI), but not Chinese from Beijing (CHB) (DeGiorgio et al., 2014). The Non-central deviation statistics NCD1 and NCD2, which measure the departure of the allele frequency spectrum from neutrality under balancing selection, showed strong evidence of balancing selection at *DMBT1* in European and African populations from the 1000 Genomes project (Bitarello et al., 2018). The beta statistic, which identifies clusters of alleles at similar frequency, identified SNPs in the 5’ region of DMBT1 to show significant balancing selection in all 1000 Genomes Project populations (Siewert and Voight, 2017). Balancing selection has been acting at this locus since before human-chimpanzee divergence, as two trans-species SNPs (rs74577795, rs79314843), a hallmark of long-term balancing selection, have been identified either side of the *DMBT1* gene (Leffler et al., 2013) (Figure 1).

**Figure 1.**
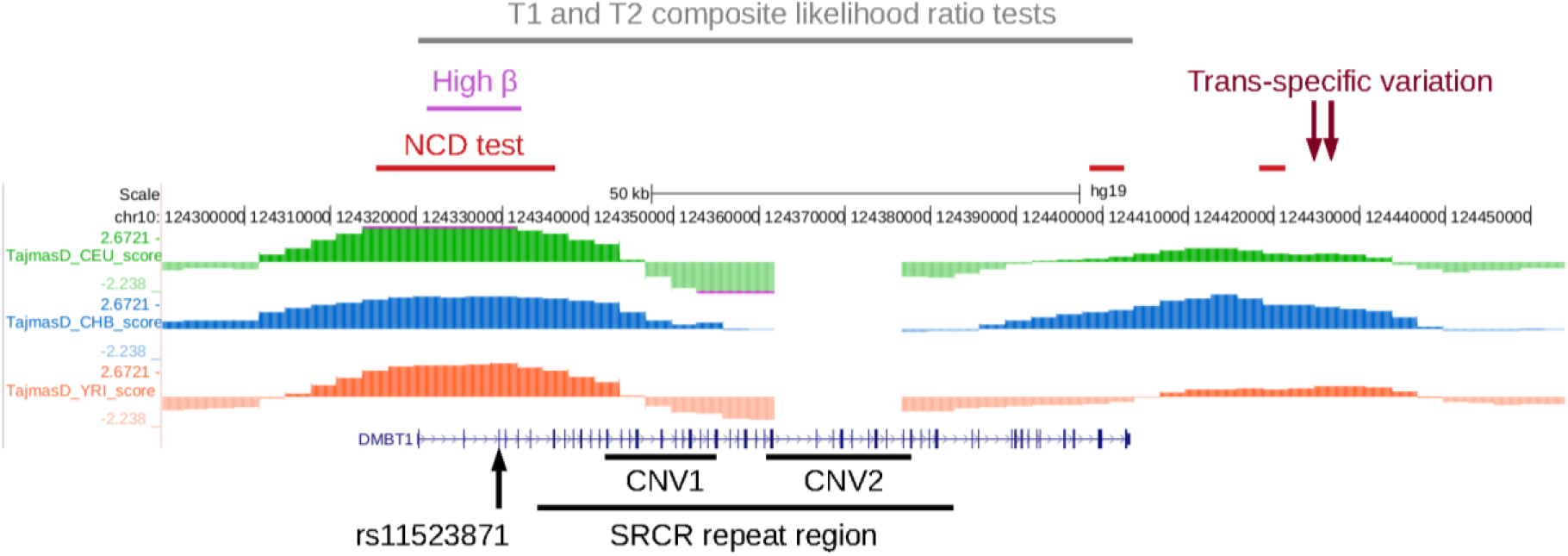
Evidence for balancing selection at the human *DMBT1* gene. The *DMBT1* gene is shown in blue, with three tracks above representing Tajima’s D measured from sequenced genomes from three populations (European-Americans from Utah (CEU) in green, Chinese from Beijing (CHB) in blue, Yoruba from Ibadan (YRI) in orange), image taken from the 1000 Genomes selection browser (https://hsb.upf.edu/). Below the *DMBT1* gene the SNP, and two CNVs which affect the copy number of the SRCR repeats, are shown, and are features analysed in this study. Above the tracks showing Tajima’s D are different sources of evidence of balancing selection, namely the beta statistic (Siewert and Voight, 2017), the NCD statistic (purple, (Bitarello et al., 2018), trans-specific variants (brown, (Leffler et al., 2013)) and composite likelihood ratio tests (DeGiorgio et al., 2014).

Genomewide selection scans for balancing selection examine patterns of SNV diversity, and so, taken together, identify regions flanking the CNV, in particular a region spanning exon 1 and the 5’ region of the gene, which shows two highly diverged haplotypes maintained at high frequencies. Whether these signals of balancing selection are independent, or are driven by linkage disequilibrium with the CNV within the *DMBT1* gene, is unclear.

Our first aim was to dissect the relationship between the evidence of balancing selection shown by the pattern of variation in regions flanking the *DMBT1* gene, and the alleles of the CNV affecting the number of SRCR domains within the *DMBT1* gene. Our second aim was to clarify the relationship between the CNV, the *DMBT1* transcript, and the different glycoprotein isoforms shown to differ functionally. Our third aim was to explore the effect of proteins from different *DMBT1* allele lengths on pattern recognition of a cariogenic and invasive *S.mutans* phenotype.

## Methods

### Source of biological material and data

The Yoruba from Ibadan, Nigeria (YRI) HapMap samples (90 DNA samples) were derived from lymphoblastoid cell lines and are available from Coriell Cell repositories.Centre de’Etude du Polymorphisme Humain (CEPH) provides a set of DNA samples derived from lymphoblastoid cell lines from European origin individuals as a three-generation pedigree that consists of 62 families with a total number of 809 individuals (Stevens et al., 2012). DNA samples with matching DMBT1 isoform data were derived from saliva collected from healthy volunteers as part of a project at the University of Umea, Sweden, with appropriate informed consent. Matching DNA and cDNA was generated from duodenal biopsies of individuals as part of a previous study (Wang et al., 1994), with local ethics approval UCLH (ref 01/0236).

H292 cell line (NCI-H292, ECACC 91091815), derived from human lung mucoepidermoid carcinomas, was supplied by the European Collection of Authenticated Cell Culture (ECACC)(Carney et al., 1985) and grown according to the supplier’s recommendation. GTex RNAseq expression data are from gtexportal.org.

### Isolation of nucleic acids

For DNA isolation, a Maxwell 16 (Promega) instrument was configured with the Standard Elution Volume (SEV) hardware and using the Maxwell 16 Cell DNA purification kit (Promega). For total RNA isolation, the SEV hardware was replaced with the Low Elution Volume (LEV) hardware and the Simply-RNA Cell kit (Promega) was used. RNA integrity was measured by electrophoresis on an Agilent Bioanalyzer 2100.

### Measurement of DMBT1 SRCR copy number

*DMBT1* copy number was determined using paralogue ratio tests (PRTs) as previously published (Polley et al., 2016a, 2016b, 2015) with long PCR used to confirm genotypes (Polley et al., 2015; Sasaki et al., 2002). Briefly, the PRT amplifies a section of DNA in a copy number variable region (test) and in a non-copy number variable region (reference) using the same primer pair. Following amplification and electrophoresis, quantification of test and reference amplicons provides a measure of the copy number of the test locus relative to the reference locus (Armour et al., 2007; Hollox, 2017). For *DMBT1*, two independent PRTs each are used to measure copy number at CNV1 and CNV2. Data analysis, and positive controls, were as previously published (Polley et al., 2015).

### SNP rs11523871 genotyping

In samples where rs11523871 genotyping data was not publicly available, it was-genotyped directly from genomic DNA using allele-specific PCR. Primers 5’TCAGTGATGGTGAATGTTTGTCA-3’ and 5’GACCTTACCTTCTGCTACAGTCGG-3’ were for the C allele; and 5’TGTGAGTGATTTATTTCGGCATTC-3’ and 5’GACCTTACCTTCTGCTACAGTCGA-3’ were for the T allele, and were included together with positive control primers from unrelated sequence 5′ATTGATTCACTTCACGGATCAAG 3′ and 5′TCTAAGAAATTCCCATGACAGGT 3′. For each sample, C-specific and T-specific reactions included primers each at a final concentration of 0.5μM, 10ng of genomic DNA, 0.2mM dNTPs and 1U Taq DNA polymerase in a standard ammonium sulfate-based PCR buffer with 1.5mM MgCl2. Standard cycling used an annealing temperature of 57°C optimised to ensure allele specificity.

### Measurement of DMBT1 gene expression using digital droplet PCR

cDNA was generated from total RNA using the Superscript III First Strand Synthesis Supermix Kit (Invitrogen). Primers and TaqMan hydrolysis probes for *RPLPO* and *UBC,* as reference genes, and *DMBT1*, were commercially available and ordered from BioRad. cDNA was digested using 50U XhoI (New England Biolabs), then 1-5ng added to a PCR mix containing 11 μl 2x ddPCR Supermix for Probe (no dUTP, BioRad), 1.1 μl 20x *DMBT1* probe assay (FAM-labelled), 1.1 μl 20x reference probe assay (HEX-labelled) in a final volume of 22 μl. After droplet generation and thermal cycling, following manufacturer’s instructions, droplets were counted by a QX200 droplet reader (Biorad) and analysed using Quantasoft (Biorad).

### DMBT1 transcript sequencing

For sequencing on a MinION (Oxford Nanopore), a sequencing library was prepared from total RNAusing the sequence-specific cDNA-PCR Sequencing kit (SQK-PCS-109, Oxford Nanopore Technologies, UK) following the manufacturer’s instructions. A primer corresponding to the 3’ end of DMBT1 (5’ [PHOS] ACTTGCCTGTCGCTCTATCTTCGCAGTTTCACCAAAATTCCTTT 3’ was used to generate cDNA from total high-quality RNA following the manufacturer’s instructions, to enrich the resulting sequence for *DMBT1* transcripts. The cDNA subsequently was amplified by PCR to enrich for full-length transcripts, following Oxford Nanopore Technology (ONT) protocols. ONT Guppy software v3.3.3 MinKNOW was used to perform live basecalling, with NanoPlot v.1.32.1 (De Coster et al., 2018) to check quality of reads. Identification of full length cDNA transcripts used Pychopper (ONT), and were mapped to chromosome 10 from the GRCh38 human genome assembly using Minimap2 v2.17 (Li, 2018). Following sorting and indexing the bam alignment files using SAMTools v1.9 (Li et al., 2009), an annotation file was generated by the spliced_bam2gff tool from the Pinfish pipeline (ONT), and visualised using the Ensembl genome browser (Yates et al., 2020).

### S. mutans strains and biotinylation

A wildtype *S. mutans* SpaP A, Cnm phenotype (strain n49) and corresponding single and double knock-out mutants were used (Esberg et al., 2017). We also generated and used a recombinant rCnm protein from the same strain. For biotinylation, bacterial cells grown overnight on blood agar plates were harvested, washed 2 times in 10mM phosphate-buffered saline pH 7.2 and resuspended in 1 ml 0.2 M NaHCO3, pH 8.3. The resultant bacterial suspension (5×10^9^ cells/ml) was mixed (10:1) with NHS-LC-biotin (Pierce, 21336P) dissolved in DMSO (1 mg/ml) and the mixture incubated with rotation for one hour at room temperature. After incubation, biotinylated bacteria were washed 3 times in PBS buffer, resuspended to a final concentration of 5×10^9^ cell/ml in adhesion buffer (50 mM KCl, 1 mM CaCl_2_, 0.1 mM MgCl_2_, 1 mM K_2_HPO_4_, pH 6.5) and stored at −20°C until use.

### Binding of S.mutans and antisera to saliva DMBT1 size isoform I-IV phenotypes

Parotid saliva (15 μL) from individuals with DMBT1 size isoform I-IV phenotypes were separated by electrophoresis on 4-15% gradient SDS-PAGE gels (BioRad) and subjected to Western blot-like binding of biotinylated bacteria. Briefly, salivary proteins were transferred to methanol treated PVDF membranes using the Trans-Blot Turbo Transfer system (BioRad). After transfer, the membrane was washed 3 times with PBS buffer with 0.05% Tween 20 (PBST), blocked in 5% milk in PBST for one hour at room temperature, washed 3 times with PBST and incubated with a suspension of biotinylated bacteria overnight at 4°C. The membrane was then washed three times (3×10 min) with PBST and bound bacteria were detected by incubating with streptavidin-POD GE Healthcare), diluted 1:10 000 in adhesion buffer, for 1 hour followed by three PBST washes (3×10 min). Bands were visualized with enhanced chemiluminescence reagent Chemidoc XRS (Biorad) and a BioRad system was used to capture the images. Western blot experiments also used specific anti-DMBT1 (mAb143 provided by D. Malamud, University of Pennsylvania), anti-PRP and anti-amylase antisera as well as pure salivary protein references to identify receptor-active saliva protein bands.

### Population genetic analysis

Statistical analysis used the statistical software R 3.6.3, implemented in RStudio v1.1.456 (https://www.R-project.org/). HGDP variation data, derived from whole genome sequencing (Almarri et al., 2020), were downloaded from the Wellcome Sanger Institute (ftp.sanger.ac.uk as vcf files) and phased using SHAPEIT2v837 (Delaneau et al., 2013). Population genetics statistics were calculated using vcftools (Danecek et al., 2011). Tajima’s D values were normalised by reporting the z value of the observed Tajima’s D value in a distribution of Tajima’s D values, measured across chromosome 10 in 16kb non-overlapping windows. Populations with fewer than three variable sites in the 16kb *DMBT1* region were excluded. Median spanning haplotype networks were calculated from phased vcf genotype data using SNiPlay (Dereeper et al., 2015).

## Results

### Two divergent haplotypes explain the footprint of balancing selection

The population genetics statistic Tajima’s D can be used to identify regions of balancing selection (Tajima, 1989). Initial inspection of Tajima’s D values across the *DMBT1* region showed a 16kb region of high value (2.67 in CEU) at the 5’end of *DMBT1* overlapping with signals of balancing selection identified using genomewide scans of other statistics (Figure 1) (Pybus et al., 2014). We calculated Tajima’s D for this 16kb region (GRCh38 chr10:122555466-122571966, Figure 1) for all populations in the Human Genome Diversity Project (supplementary table 1). As expected, a high Tajima’s D value was shown across almost all European populations and African populations, with large differences in values sometimes observed between closely-located populations, particularly in Asia.

We examined the haplotype structure of the 16kb region in the French population and the Yoruba population, both showing a high positive Tajima’s D in sequencing data from the Human Genome Diversity Panel. Two highly diverged haplotypes were responsible for the signal of balancing selection in this and other populations (supplementary figure 1. Therefore, the allelic status at many SNPs in this region can distinguish the two major haplotype clades (Supplementary figure 1), and we chose SNP (rs11523871) to act as a proxy for the two haplotypes under balancing selection.

### Copy number variation of DMBT1 SRCR domains is associated with the two haplotype clades

Our hypothesis was that the high frequency of the divergent haplotypes was driven by linkage of each haplotype to distinct spectra of copy number variants in the *DMBT1* gene. To explore whether the copy number of the tandemly repeated SRCR domains is responsible for the signal of balancing selection that the haplotypes, we phased individual SRCR domain copy numbers to haplotypes by observing segregation in families from the CEPH cohort. rs11523871-C haplotypes carry fewer SRCR repeats compared to rs11523871-A haplotypes (p=3.282×10^-7^, n=263, Wilcoxon rank sum test with continuity correction), with the relationship mostly explained by a higher frequency of the short 8 SRCR domain allele on the rs11523871-C haplotype (Figure 2a). We also phased individual SRCR domain copy numbers to haplotypes in a cohort of Yoruba individuals from Ibadan Nigeria (YRI), using parent-offspring trio information and the software CNVice (Zuccherato et al., 2017). Again, we found that rs11523871-C haplotypes carry fewer SRCR domain repeats compared to rs11523871-A haplotypes (p=5.5×10^-4^, n=116, Wilcoxon rank sum test with continuity correction), but this association is due primarily to a high frequency of the 11 allele on the rs11523871-C haplotypes (Figure 2b). In summary shows, association between SNP alleles and SRCR domain copy numbers is population-specific, with the rs11523871-C haplotype carrying, on average, a shorter SRCR domain than the rs11523871-A haplotype.

**Figure 2.**
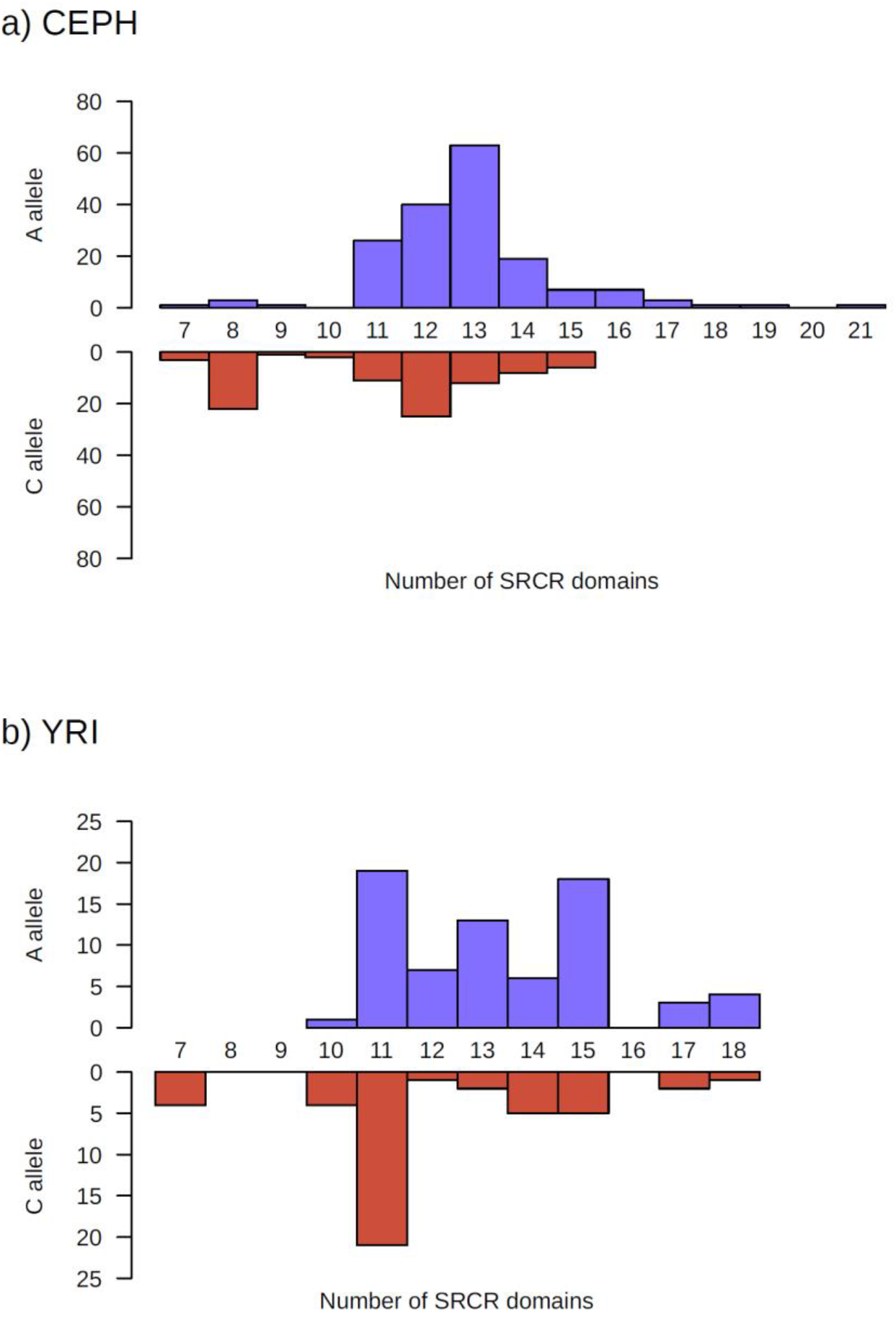
Association of rs11523871 and *DMBT1* SRCR repeat copy number. For two populations, CEPH (a) and YRI (b), the distributions of *DMBT1* SRCR repeat domain copy numbers associated with the rs11523871-A allele (blue, above the x-axis) and rs11523871-C (red, below the x axis) are shown. The y axis shows the number of observations in the two samples (CEPH n=263, YRI n=116).

We also explored other explanations for the high frequency of divergent haplotypes. Firstly, we explored whether any of the variants in the divergent haplotypes were likely to have a direct effect on the DMBT1 protein. There are two common SNPs that alter an amino acid, rs11523871 (Thr42Pro) and rs3013236 (Leu54Ser), both outside the functional SRCR domains. Analysis using Polyphen and SIFT through Ensembl’s Variant Effect Predictor (McLaren et al., 2016) reported these variants as benign and tolerated, respectively, arguing against these SNPs causing a functional difference and therefore theory are unlikely to be responsible for the signal of balancing selection.

If the genotype of one of the variants within the divergent haplotype were to affect expression levels of the gene, it is possible that this could explain the pattern of balancing selection observed. Analysis of GTex data confirmed the established tissue expression pattern, with high expression levels in lung, small intestine, colon and minor salivary gland (Figure 3a). Of these tissues, only in the colon does *DMBT1* expression levels show a relationship with genotype at rs11523871 (p=1.7×10^-19^, n=368), with a linear increase in expression with each C allele, with the CC genotype showing on average the highest expression (Figure 3b). Given the lack of evidence in other tissues, and the uncertainty surrounding measuring the abundance of transcripts containing tandem repeats using short-read RNAseq, we tested the relationship between rs11523871 genotype and expression level in the duodenum from 41 individuals using digital droplet PCR (Figure 3c). This shows variation in expression levels (ANOVA, p=0.027), and weak support for a linear relationship with the C allele (p=0.047, *UBC* as reference, p=0.09, *RPLP0* as reference, two-tailed F-test). Taken together, there is no evidence of a heterozygous effect on expression level, but a relationship between allele C, acting as a proxy for one haplotype at the balancing selection locus at rs11523871 and high expression.

**Figure 3.**
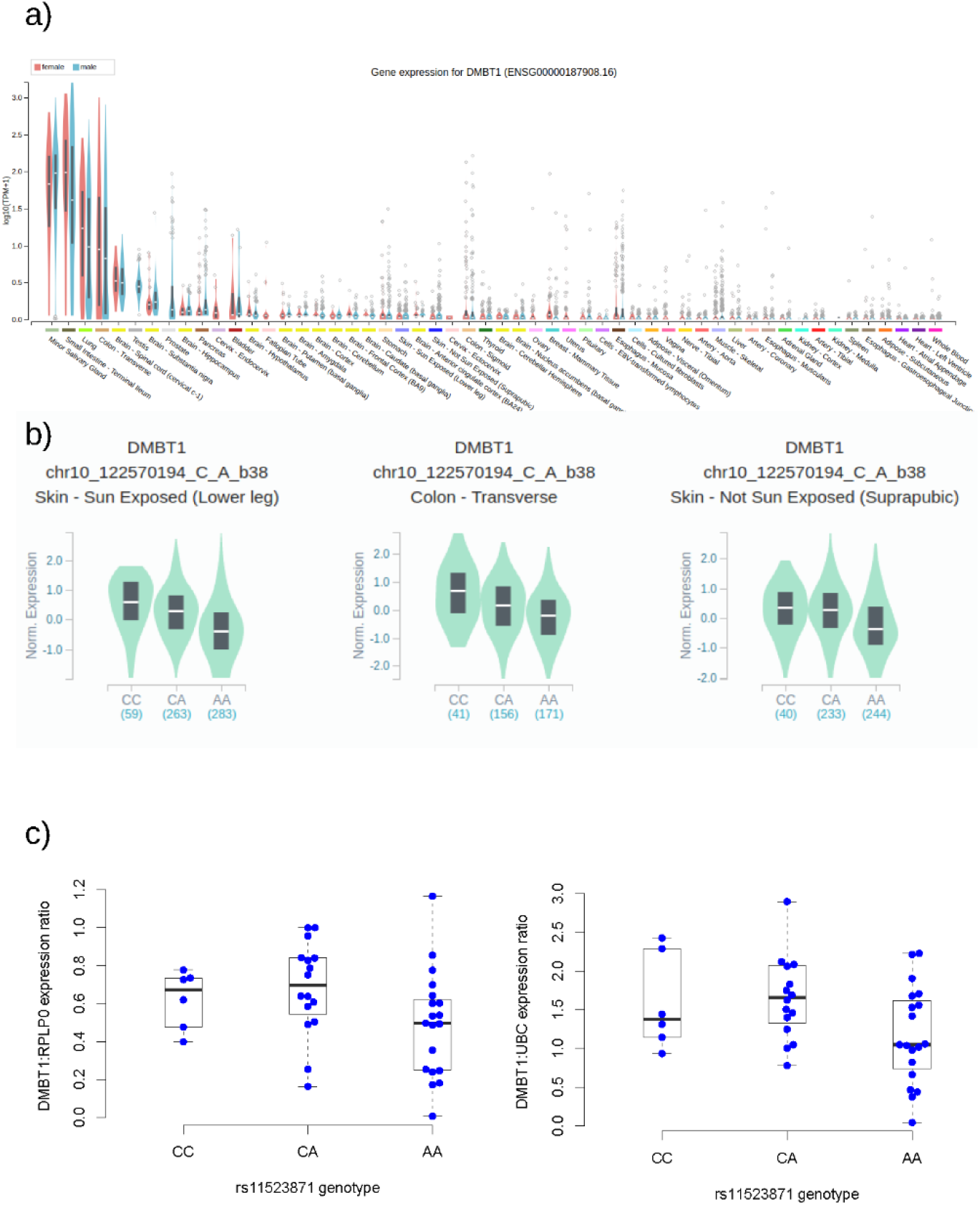
*DMBT1* gene expression and rs11523871 genotype. a) Tissue expression of *DMBT1* across 54 tissues, ordered by mean expression level, from RNAseq data. Data and image from the GTEx Portal Locus Browser v.8 (https://gtexportal.org). b) Violin plots show rs11523871 genotype and expression level for the three tissues showing a statistically significant relationship. c) Boxplots showing rs11523871 genotype and expression level in duodenum from 41 healthy patients, normalised against two different housekeeping genes. Left boxplot shows data normalised to *RPLP0* expression, right boxplot shows data normalised to *UBC* expression.

### Copy number variation of DMBT1 SRCR domains is associated with DMBT1 protein size variation in saliva

It is known that cloned *DMBT1* transcripts of different lengths generate proteins of different lengths, but the basis of the size variation in DMBT1 protein isoforms isolated directly from saliva, for example, is unclear. To address this, we determined SRCR domain diploid copy number on eight DNA samples from individuals that had previously been assigned as different protein isoforms (I-IV) by western blot of saliva (Eriksson et al., 2007) (Table 1). Even with a small sample size, a strong linear relationship between SRCR diploid copy number and DMBT1 protein isoforms was present (linear regression, p=0.005, Kendall’s rank correlation p=0.004), explaining most of the variation in protein size (r^2^=0.75).

**Table 1.** DMBT1 protein isoforms and SRCR repeat diploid copy number.

| Sample | CNV1<br>diploid<br>copy<br>number | CNV2<br>diploid<br>copy<br>number | SRCR<br>repeat<br>diploid<br>copy<br>number | Inferred<br>Secretor<br>status | DMBT1<br>(gp340)<br>protein<br>isoform | Approximate<br>isoform<br>size*<br>(kDa) | *S. mutans*<br>binding<br>activity<br>** |
| --- | --- | --- | --- | --- | --- | --- | --- |
| 1 | 2 | 7 | 27 | + | III | 389 | + |
| 2 | 2 | 4 | 24 | - | I | 345 | - |
| 3 | 2 | 2 | 22 | + | I | 345 | - |
| 4 | 2 | 6 | 26 | + | II | 375 | + |
| 5 | 2 | 4 | 24 | + | II | 375 | + |
| 6 | 1 | 4 | 20 | - | IV | 287,345 | - |
| 7 | 2 | 5 | 25 | + | II | 375 | + |
| 8 | 2 | 9 | 29 | + | III | 389 | + |

\*Protein isoform data from (Eriksson et al., 2007)
\*\* see Figure 5

We also explored the possibility of alternative splicing in generating DMBT1 variants. Genetic variation in SRCR domain copy number must be reflected in the transcript and the protein if balancing selection operating on the function of the protein is reflected in genetic variation. Long (8 kb, 13 SRCR domains) and short (6kb 8 SRCR domains) transcripts have been cloned from cDNA and studied extensively, yet different transcript lengths have previously often been ascribed to alternative splicing between tissues rather than genetically-encoded polymorphic variation. To help resolve this issue, we took a H292 lung cell line model, where *DMBT1* is strongly expressed, and used long single molecule sequencing to determine the extent to which the copy number of SRCR domain repeats in *DMBT1* transcripts reflects the copy number of SRCR domain repeats in the *DMBT1* gene (Figure 4). Using PRT and long PCR, we showed that the H292 cell line was homozygous for 11 SRCR domain repeats. Five transcripts with mappable 3’ and 5’ ends were identified, of which four showed 11 SRCR domain repeats and one showed 10. This suggests that, for this cell line model at least, alternative splicing can generate alternative numbers of SRCR domain repeats but plays a relatively minor role, with the length of most transcripts matching the genetically encoded number of SRCR domains.

**Figure 4.**
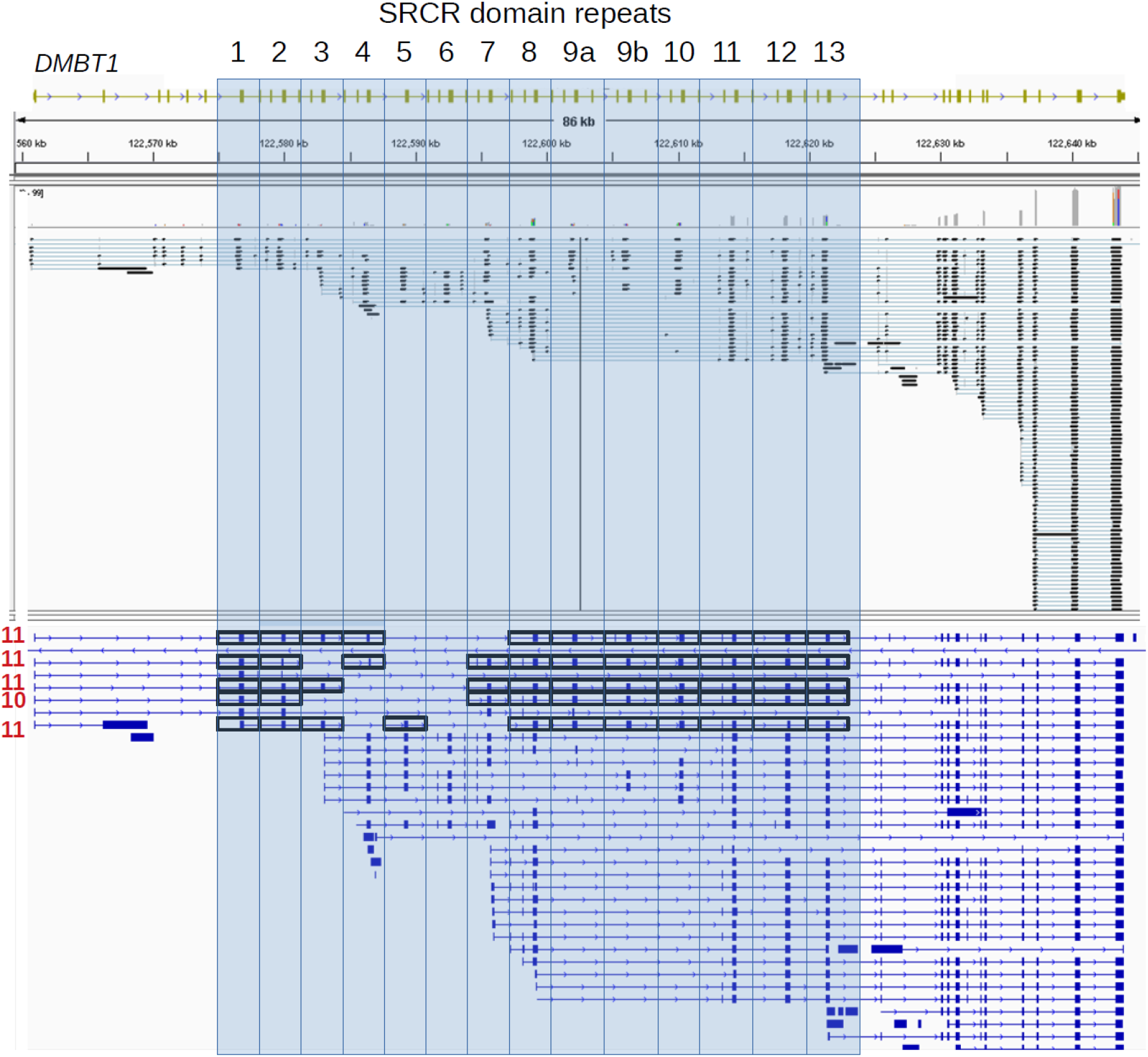
Identification of transcripts spanning the SRCR repeats in *DMBT1*. The *DMBT1* allele from the genome assembly, with 14 tandemly-arranged SRCR repeats highlighted in blue, is shown at the top of the figure, with GRCh38 coordinates as a scale immediately underneath. The sequence alignment of Nanopore single molecule sequencing reads mapping to *DMBT1* are shown, with at the bottom, the genome features format file (GFF) derived from the sequence alignments. In the GFF image, black boxes indicate complete SRCR repeats, with the total number of tandemly-repeated SRCR repeats for that transcript highlighted in red on the left of the particular transcript.

### A short DMBT1 SRCR domain allele shows loss of pattern recognition of a cariogenic and invasive S. mutans phenotype

We next explored the ability of DMBT1 protein size isoforms I-IV to bind a *S. mutans* SpaP A, Cnm phenotype using a Western blot-like assay and salivas with known DMBT1 size isoforms I-IV (Eriksson et al., 2007). Both spaP and Cnm are adhesins expressed on the surface of *Streptococcus mutans* thought to mediate interactions with salivary DMBT1 (Brady et al., 2010, 1992; Esberg et al., 2017; Loimaranta et al., 2005). DMBT1 size isoforms II and III (carrying longer alleles) mediated distinct binding, while isoforms I and IV (carrying shorter alleles) did not bind the *S. mutans* ligand (Figure 5, Table 1). Both silver staining and Western blot with anti-DMBT1 antisera verified the amount of DMBT1 in the salivas (Figure 5). Binding to saliva of single (spaP A-or Cnm-) and double (spaP A-, Cnm-) knock-out mutants of the wild-type strain (spaP A, Cnm), and recombinant rCnm adhesin, to saliva indicated that binding to DMBT1 was mediated mainly by the Cnm adhesin (Figure 5). The weak residual activity upon binding of the double mutant (spaP A-, Cnm-) may reflect other adhesins other than Cnm or spaP (Supplementary figure 2).

**Figure 5.**
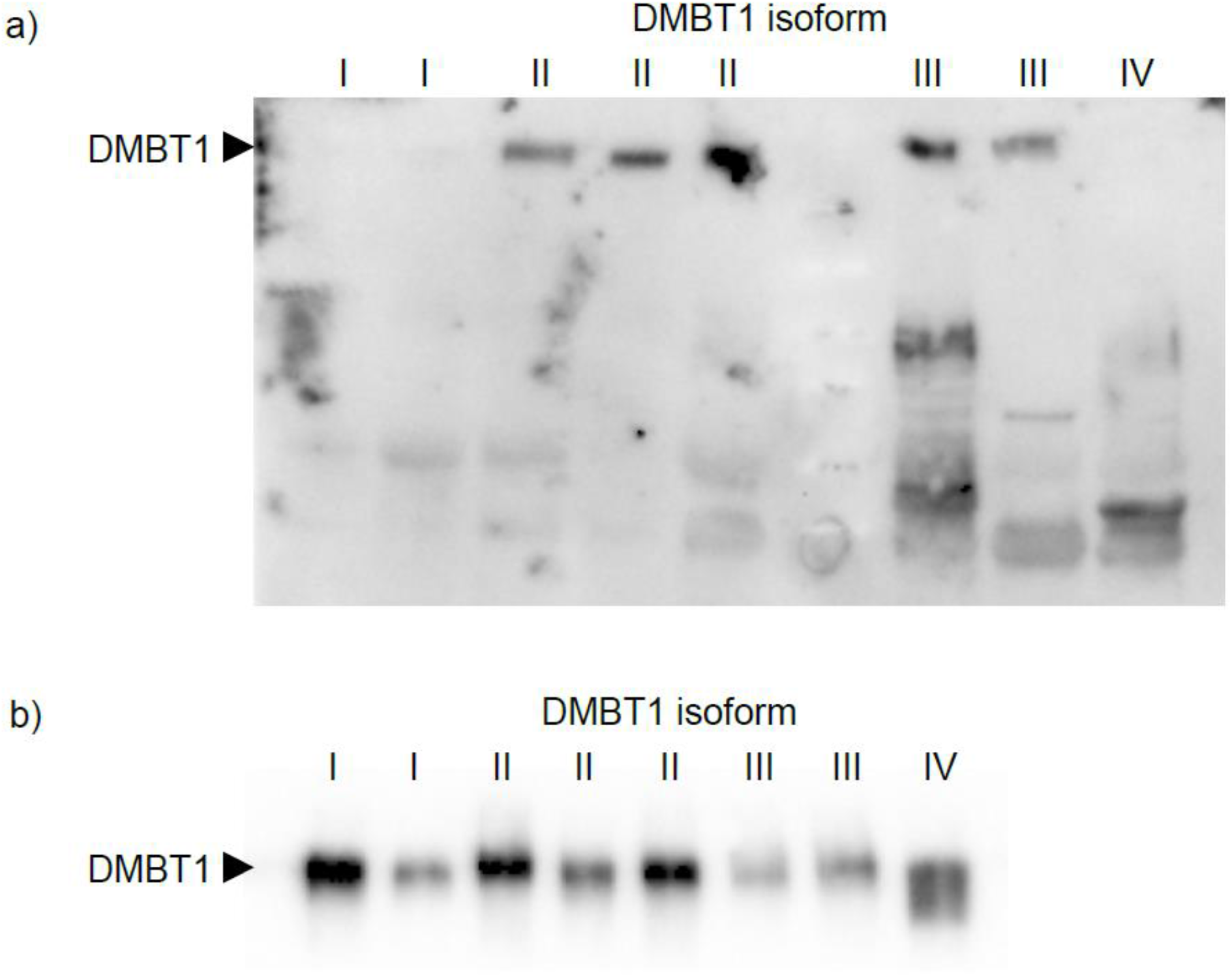
Differential binding of *S.mutans* by DMBT1 isoforms in saliva. Overlay of individual saliva phenotypes with DMBT1 size isoforms I-IV with A) a biotinylated *S. mutans* SpaP A, Cnm strain and B) with DMBT1-specific antibodies. The positions of DMBT1, mono- and dimer amylase, and acidic PRP co-receptors are marked by arrows.

Binding of *S. mutans* occurred also to mono and dimeric amylase (50 and 100 kDa) and PRP (43 and 37 kDa) protein bands, although with large individual variation. Recombinant Cnm bound DMBT1 and amylase but not PRP (Supplementary figure 2).

### Variation in Secretor-dependent glycosylation may contribute to DMBT1 protein isoform variation

Previous reports have suggested that differential glycosylation may explain the variation in DMBT1 isoform size observed, and indeed DMBT1 shows complex glycosylation patterns, including glycosylation by the product of the *FUT2* gene, where a null allele is responsible for the Secretor negative blood group, and shows evidence of balancing selection itself (Koda et al. 2000; Soejima et al. 2007; Ferrer-Admetlla et al. 2009). We genotyped the *FUT2* gene to infer Se status, and found 2 Se- and 6 Se+ individuals. Incorporating this into our linear model improved the fit of the model to the data (r^2^=0.85), although secretor status was not a statistically significant predictor (p=0.07). It is clear that SRCR domain copy number is primarily responsible for the different protein isoforms of DMBT1, although Se status, and other glycosylation, may contribute.

## Discussion

In this study, we have shown that the strong signature of balancing selection spanning the 5’ region of the *DMBT1* gene is most likely to be explained by tight linkage of particular SNP haplotypes at this region to particular SRCR domain repeat copy numbers within the *DMBT1* gene. In particular, the signal of balancing selection across populations is driven by one haplotype carrying a short SRCR domain repeat copy number allele and another carrying a longer SRCR domain repeat copy number allele. We also show that SRCR domain repeat copy number is the primary determinant of both transcript length and protein isoform size in vivo, and that the short protein isoform shows complete loss of pattern recognition or binding to a cariogenic and invasive *S.mutans* phenotype.

The finding of the short protein isoform showing complete loss of binding of the collagen-binding Cnm *S.mutans* phenotype associated with caries and invasive,systemic conditions is interesting for several reasons. The fact that previous studies have shown a 30%-40% reduction of binding of other *S.mutans* types and other bacteria suggests that the longer DMBT1 isoform may give optimal protection against the Cnm and other collagen-binding phenotypes with the potential to cross the basal membrane barrier and infect the extracellular matrix. However, our data suggest that other co-receptors, besides DMBT1, may contribute to the partial reduction of *S.mutans* binding when whole saliva is analysed. Although the number of individuals in our study are too few for firm conclusions, our present findings are consistent with previous findings (Eriksson et al., 2007) in suggesting that the short DMBT1 isoform may lack *FUT2*-dependent fucose receptors for many microbial ligands. Studies showing a role for DMBT1 genetic and protein variation in individuals with high risk of caries, and different causal subtypes, indicate the clinical significance of DMBT1 (Jonasson et al., 2007; Esberg et al., 2017; Strömberg et al., 2017).

Taken together, our work and previous work showing that DMBT1 molecules with fewer SRCR repeat domains are less effective at binding bacteria provides a link between the observation of balancing selection in the genome and the functional basis for that balancing selection (Bikker et al., 2017). The exact nature of the balancing selection remains unclear. For example, could it be due to overdominance, where heterozygotes with a long and a short *DMBT1* allele are favoured. An alternative mechanism could be fluctuating selection, where a change in the environment across space or time alters the selective advantages of different alleles. In support of this, the observation that the Tajima’s D value of the linked region can vary wildly between closely-located populations (Table S1) with more subtle differences in the SRCR copy number (Polley et al., 2015). A possible scenario is that alleles reducing tethering to teeth are favoured in populations with cereal-rich diets, alleles with enhanced binding to bacteria are favoured during outbreaks of diarrheal or lung disease, and alleles with improved viral binding signalling to complement and favoured during viral infectious disease epidemics.

Any or all of these explanations are plausible, and while it is possible to determine relative importance of the different alleles on these different functions, the actual selection pressures that have acted on *DMBT1* and are responsible for the variation that we say are likely to be very difficult to determine.

Finding explanations for patterns of balancing selection in the human genome can have medical importance, as it can identify alleles that have a functional effect in a disease process. Examples include loci under balancing selection for malaria, where GWAS have supported the association. Loci under balancing selection may identify further loci invisible to current GWAS either because of limitations in measuring phenotypes or the transient complex nature of other phenotypes, such as infectious disease. It is notable that, as yet, there is no convincing association of variation at *DMBT1* and disease - there are no GWAS-based associations in the region of balancing selection, and early suggestions that the number of SRCR repeat domains was associated with Crohn’s disease (CD) (Renner et al., 2007) have not been replicated (Polley et al., 2016b), although there is some evidence of an association of CD with rs2981804 in a candidate gene study (Diegelmann et al., 2013), and recent evidence suggests a genetic and functional link between SRCR copy number and the number of urinary tract infections in children with vesicoureteral reflux (Hains et al., 2021). However, most links between *DMBT1* and diseases such as lung disease rely entirely on evidence from functional studies (Hartshorn et al., 2006; Müller et al., 2015, 2008).

In conclusion, our work supports the concept that selection acts on a highly-mutable SRCR domain-encoding tandem repeat in the *DMBT1* gene in humans, at least in part, by differential binding properties of the molecule to co-receptors on the tooth surface and diverse bacteria on mucosal surfaces. The expression of DMBT1 in saliva, lung, intestinal surfaces, amniotic fluid, and as a major protein (4-10%) in the first meconium stool of newborns, may suggest an important role in the host’s interaction with both pathogenic and commensal bacteria. Understanding the consequences of variation of this molecule in the mucosal innate immune response to infection will be an important aspect of understanding the role of *DMBT1* in health and disease.

## Acknowledgements

Thanks to Rachael Madison for technical support. Access to a Biorad QX200 digital droplet generator and reader was through the NUCLEUS Genomics service at the University of Leicester. We thank Prof Dallas Swallow for access to intestinal DNA and cDNA samples. AA was supported by the Saudi Arabian Ministry of Health, and by a PhD studentship grant from the Saudi Arabian Cultural Bureau, London.

## Supplementary figures

**Supplementary Figure 1.**
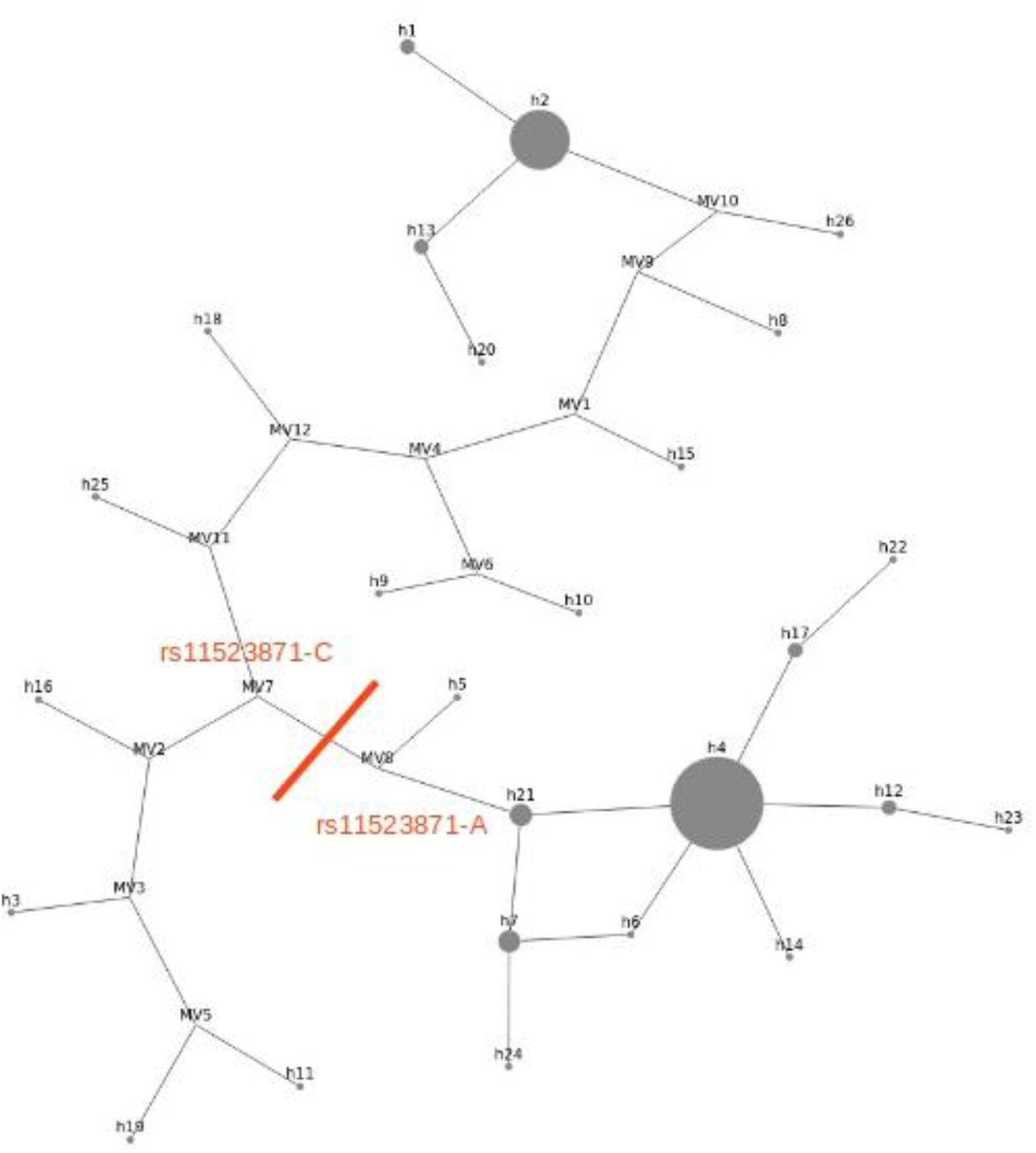
Haplotype network of *DMBT1* 16kb region in the French population. Haplotype median joining network of GRCh38 chr10:122555466-122571966 showing balancing selection and high Tajima’s D values. The line indicates the boundary between haplotypes carrying C and A at rs11523871 and A, h indicates observed haplotypes, with size of circle representing number of observations in the sample, MV indicates internal nodes with no observed representative haplotype.

**Supplementary Figure 2.**
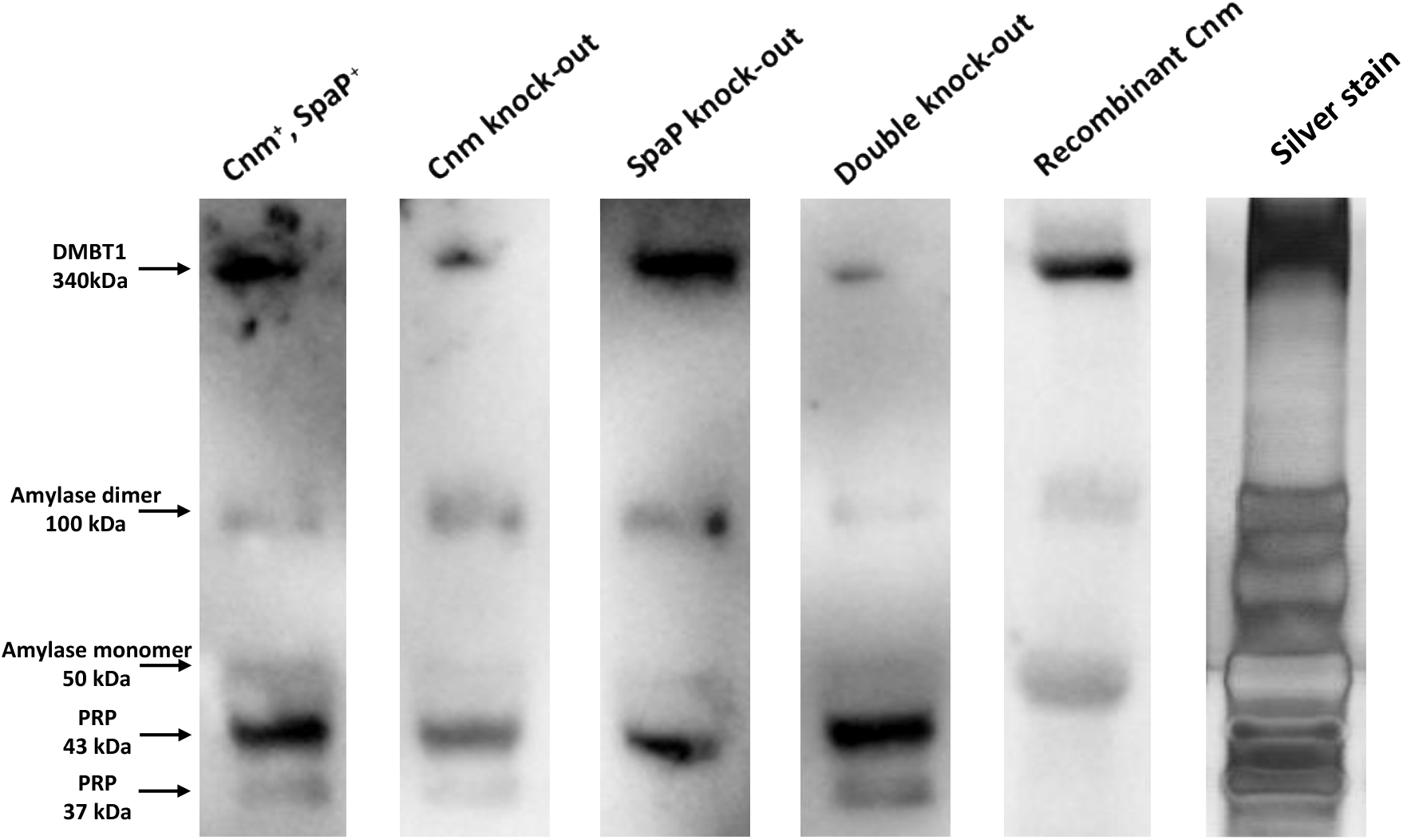
Binding of *S. mutans* phenotypes and recombinant rCnm to DMBT1 and co-receptors in saliva. Saliva proteins blotted to a membrane after separation of parotid saliva were overlaid with wildtype (spaP A+, cnm+), single (spaP A- or cnm-) and double (spaP A-, cnm-) knock-out mutants and rCnm. The wildtype strain and Cnm+, spaP-mutant and rCnm protein showed equal binding to DMBT1, while the spaP A+, Cnm- and spaP A-, cnm-mutants showed reduced but residual activity. Both strains, mutants and rCnm bound to mono- and dimer amylase but strains/mutants and not rCnm to the acidic PRP protein bands, suggesting a plausible unspecific binding to the major saliva amylase components but potentially specific features of the acidic PRP co-receptors. The amylase and PRP components are marked by arrows.

## Supplementary table

**Supplementary Table 1.** Normalised values for DMBT1 Tajima’s D in HGDP populations GRCh38 chr10:122555466-122571966.

| Population | z_value_TajimaD |
| --- | --- |
| Adygei | 2.76 |
| Balochi | 1.49 |
| Bantu_NE | 3.22 |
| Bantu_S | 3.75 |
| Bedouin | 3.27 |
| Biaka | 3.44 |
| Brahui | 2.00 |
| Burushi | 1.08 |
| Cambodian | 2.36 |
| Dai | 0.64 |
| Daur | -0.27 |
| Druze | 1.73 |
| French | 3.36 |
| French_Basque | 3.63 |
| Han | 1.19 |
| Hazara | 0.75 |
| Hezhen | -1.25 |
| Japanese | 0.95 |
| Kalash | -0.01 |
| Lahu | -0.07 |
| Makrani | 2.15 |
| Mandenka | 4.02 |
| Maya | -1.41 |
| Mbuti | 1.90 |
| Miao zu | 2.13 |
| Mongola | -0.32 |
| Mozabite | 4.21 |
| NAN_Melanesian | -0.94 |
| Naxi | -0.50 |
| North_Italian | 0.55 |
| Orcadian | 2.33 |
| Oroqen | -1.15 |
| Palestinian | 3.11 |
| Papuan | -1.78 |
| Pathan | 2.16 |
| Pima | -1.95 |
| Russian | 2.77 |
| San | 1.60 |
| Sardinian | 3.41 |
| She | 2.45 |
| Sindhi | 0.96 |
| Tu | 1.00 |
| Tujia | 2.28 |
| Tuscan | 1.90 |
| Uygur | 1.77 |
| Xibo | -0.41 |
| Yakut | 0.51 |
| Yizu | 0.94 |
| Yoruba | 4.29 |

